# Multi-omic modeling of antidepressant response implicates dynamic immune and inflammatory changes in individuals who respond to treatment

**DOI:** 10.1101/2022.10.07.511374

**Authors:** Shih-Chieh Fuh, Laura M. Fiori, Gustavo Turecki, Corina Nagy, Yue Li

**Affiliations:** School of Computer Science, McGill University, 3480 Rue University, Montréal, QC H3A 2A7, Canada; Department of Psychiatry, McGill Group for Suicide Studies, Douglas Mental Health University, Montreal, QC H4H 1R3, Canada

## Abstract

Major depressive disorder (MDD) is a leading cause of disability worldwide, and is commonly treated with antidepressant drugs (AD). Although effective, many patients fail to respond to AD treatment, and accordingly identifying factors that can predict AD response would greatly improve treatment outcomes. In this study, we developed a machine learning tool to integrate multi-omic datasets (gene expression, DNA methylation, and genotyping) to identify biomarker profiles associated with AD response in a cohort of individuals with MDD. To address this rich multi-omic dataset with high dimensional features, we developed integrative Geneset-Embedded non-negative Matrix factorization (iGEM), a non-negative matrix factorization (NMF) based model, supplemented with auxiliary information regarding genesets and gene-methylation relationships. Using our model, we identified a number of meta-phenotypes which were related to AD response. By integrating geneset information into the model, we were able to relate these meta-phenotypes to biological processes, including immune and inflammatory functions. This represents both biomarkers to predict response, as well as potential new treatment targets. Our method is applicable to other diseases with multi-omic data, and the software is open source and available on Github (https://github.com/li-lab-mcgill/iGEM).

## Introduction

Major depressive disorder (MDD) is a chronic and debilitating illness that affects 300 million people worldwide^1^. Its core symptoms are depressed mood and lack of interest, altered appetite and sleep, difficulty concentrating, and suicidal thoughts^2^. Between one-to two-thirds of MDD patients do not respond to their first line of antidepressant (AD) treatment, and up to one-third of patients fail to respond after multiple courses of AD treatment^3^. As such, identifying factors which can predict AD response could greatly improve treatment outcomes.

A variety of clinical and sociodemographic variables have been investigated as predictors of response, such as diagnostic symptomology, age, socioeconomic variables, and BMI^4, 5^. However, the use of molecular measures to predict response has become increasingly important, in part as they have not only the potential to predict response, but also to reveal underlying biological mechanisms^6-9^. Although the brain represents the most relevant tissue for identifying biomarkers of AD response, due to its lack of accessibility, the majority of studies rely on the investigation of peripheral markers^10^. A number of high-throughput studies have been performed in the blood, including those investigating gene expression, DNA methylation, genetic variation, and metabolic profiling, and have identified a number of factors related to AD response^11-17^. While promising, many of these findings have yet to be replicated, nor is it clear how findings from these different studies may be related. Moreover, it has become abundantly clear from genome-wide analyses that both MDD and AD response is highly polygenic and pleiotropic, each with small effect sizes. This creates challenges when using genome-wide approaches, particularly with smaller sample sizes, which may lack sufficient power to detect relevant, but small, differences. Additionally, it is likely that the molecular factors that predict AD response involve both static (ie, genetic variation) and dynamic effects (ie, gene expression). Accordingly, the integration of multi-omic datasets may provide a greater capacity to predict response. While previous studies have implemented multi-omic approaches to study AD response and MDD, for example, by combining genotype and metabolite data^18^, our study increases the dimensions by including three levels of molecular profiling and clinical data.

In the present study, we integrated gene expression, DNA methylation, genotype variation, and clinical questionnaire scores from a cohort of individuals with MDD who were treated with AD for 8 weeks, allowing us to generate a composite biomarker of AD response. To investigate this multi-omic dataset, we applied non-negative matrix factorization (NMF). Originally developed by Lee et. al.^19^, NMF has been applied to the field of bioinformatics in various contexts, including cancer classification and feature clustering^20, 21^. It is a powerful framework for analysis of multi-omic datasets because it can distill latent factors from the data, which we often refer to as *topics* below. In this context, NMF can produce patient loading scores and feature basis scores, representing the relative importance or contribution of the patients and features (such as the genes) with respect to each topic. As a result, we can treat each topic as a *metagene* and perform hypothesis testing over these *metagenes*, which are much fewer than the total number of genes, thereby reducing the number of statistical tests being performed.

These characteristics of NMF make it suitable for multi-omic dataset analyses, and several researchers have applied NMF towards investigating MDD, with promising results. Shao et. al. used a cluster-driver NMF to discover common and distinct brain structural connectivity patterns between individuals with schizophrenia and MDD^21^. In addition, a recent pilot study by Chen et. al. applied NMF on the 30-item Inventory of Depressive Symptomatology-Clinician Rating (IDS-C30) to derive subtypes of treatment-resistant depression (TRD) and identified biomarkers with differential expression between the subtypes, including several immune and inflammation regulators such as IL-1β, IL-6, and TNF-α^22^. However, NMF alone does not directly reveal pathway or geneset information and often requires *post-hoc* geneset enrichment analysis to interpret the topics.

In this study, we present integrative Geneset-Embedded non-negative Matrix factorization (iGEM) to computationally address the above-mentioned challenges. Briefly, iGEM is a joint NMF model with sparsity constraint and biological network regularization. To guide topic learning, iGEM exploits known geneset information by further factorizing the feature factor score matrix into topics-by-genesets and genesets-by-genes. In this study, we applied iGEM to multi-omic data to derive shared factor scores of genes, genesets, and methylation sites for the relative importance in each topic. With the factor scores, we performed additional analyses to incorporate genotype and clinical feature information, to better understand the relationships between molecular processes and MDD, and how they relate to AD response.

## Methods

### Data

#### Patient cohort

Participants were recruited from an outpatient clinic at the Douglas Mental Health University Institute in Montréal, Canada, between February 2012 and October 2016. All individuals were diagnosed with major depressive disorder, and had an active episode when recruited. Exclusion criteria included comorbidity with other major psychiatric disorders, bipolar disorder, alcohol or substance abuse over the last 6 months, and a concurrent physical illness. All subjects were free of psychotropic medication at baseline (T0). Patients were treated for 8 weeks (T8) with either i) Desvenlafaxine, a serotonin–norepinephrine reuptake inhibitor, started at 50 mg die and increased to 100 mg if needed; or ii) Escitalopram, started at 10mg die and increased to 20mg if needed. All subjects were assessed at baseline and after 8 weeks (T8) using the Montgomery–Åsberg Depression Rating Scale (MADRS), and were classified as responders (RES) (N=83) or non-responders (NRES) (N=79) based on a ≥ 50% reduction in scores from baseline^23^. This study was approved by our local institutional review board, and all participants provided written informed consent. Only patients with data available for gene expression, methylation, genotype, and MADRS response at both week 0 and week 8 were included, giving a final subject count of 61 RES and 50 NRES (,**Supp. Table 1**).

#### Blood samples

Peripheral blood samples were collected at baseline, and tubes were frozen using a sequential freezing process. DNA was extracted from PAXgene DNA tubes (PreAnalytix) using a modified version of the Qiagen FlexiGene DNA kit, and was stored at −20°C. Whole blood for RNA was collected in EDTA tubes and filtered using LeukoLOCK filters (Life Technologies). Total RNA was extracted using a modified version of the LeukoLOCK Total RNA Isolation System protocol, and included DNase treatment to remove genomic DNA. RNA quality was assessed using the Agilent 2200 Tapestation, and only samples with RNA Integrity Number (RIN) ≥ 6.0 were used.

#### Gene expression

RNA sequencing was performed using RNA extracted at baseline for 150 samples (79 responders, 71 non-responders). All libraries were prepared using the Illumina TruSeq mRNA stranded protocol following the manufacturer’s instructions. Samples were sequenced at the McGill University and Genome Quebec Innovation Centre (Montreal, Canada) using the Illumina HiSeq4000 with 75nt paired-end reads. FASTXToolkit and Trimmomatic were used for quality and adapter trimming, respectively^24, 25^. Tophat2, using bowtie2, was used to align the cleaned reads to the reference genome (GRCh38)^26, 27^. Reads that lost their mates through the cleaning process were aligned independently from the reads that still had pairs. Quantification on each gene’s expression was estimated using HTSeq-count and a reference transcript annotation from ENSEMBL^28^. Counts for the paired and orphaned reads for each sample were added to each other. Normalization was conducted on the resulting gene matrix using DESeq2^29^. Counts thus normalized were log 2 transformed and corrected for age, sex, and RIN using the removeBatchEffect function of limma^30^. Genes with counts lower than 30 were filtered. A total of 58126 genes passed the filter and were then applied to the subsequent differential analysis.

#### Methylation

DNA extracted at baseline for 147 samples (71 responders, 76 non-responders) was bisulfite treated using the EZ DNA Methylation-Gold Kit (Zymo Research), and hybridized to the Infinium MethylationEPIC Beadchip (Illumina). The Infinium MethylationEPIC Beadchip was used to assess genome-wide DNA methylation (Illumina, US). After accounting for attrition rates, and DNA sample quality control, pre-processing and analysis of raw microarray data for the remaining samples was conducted within R (version 3.4) predominantly using the Chip Analysis Methylation Pipeline (ChAMP) Bioconductor package, which utilizes many elements of minfi^31, 32^. Sample methylation signal QC was assessed by plotting log median methylated and unmethylated signals. Samples were removed if they failed to cluster with others or if they exhibited lower median intensities in either signal channel. Probes with low signal detection relative to control probes, probes with < 3 beads in > 5% of samples, cross reactive probes, non-CpG probes, sex chromosome probes, and probes that hybridize to single nucleotide polymorphism sites (SNPs) were removed. Beta values were calculated as the ratio of methylated signal to the sum of unmethylated and methylated signals at each CpG site, and subsequently normalized. Log2 transformed beta values were used for the remainder of pre-processing steps as recommended by Du et al., but reported as beta values^33^. Technical batches and covariates were detected using single value decomposition analysis. Detected and known batch effects were corrected for prior to differential methylation analysis. The CpG site annotations are based on the chip manifest (the manifest uses information from the UCSC)^26^. CpG sites are annotated to a gene if they are in the body or less than 1500bp upstream of the transcription start site (TSS). Technical batches and covariates were detected using single value decomposition analysis. Detected and known batch effects were corrected for prior to differential methylation analysis. We used DNA methylation beta values from 673039 methylation sites for the differential analysis.

#### Genotype

Genotyping was performed for 157 subjects (76 responders, 81 non-responders) using the Infinium PsychArray Beadchip (Illumina). The genotype information from a total of 598131 SNPs were then filtered through PLINK^34^. The SNP variations were encoded as 0, 1, and 2.

#### Depression scores

We used both total MADRS scores, as well as scores from individual items. Scores for each item range between 0 to 6, with higher scores indicating more severe symptoms. For use in the model, we defined the total score as the difference between total scores at week 0 and week 8, divided by of the total score at week 0. The MADRS questionnaire includes 10 items covering sadness (reported sadness and apparent sadness), negative thoughts (pessimistic and suicidal), detachment (concentration, lassitude, feeling), and neurovegetative symptoms (inner tension, reduced sleep, and reduced appetite)^35^. Sadness includes reported sadness and apparent sadness. Negative thoughts include pessimistic thoughts and suicidal thoughts. Inner tension, reduced sleep, and reduced appetite belong to the neurovegetative symptoms. Detachment covers concentration difficulties, lassitude, and inability to feel.

#### Genesets

An integrated human geneset dataset with a total of 4337 Genesets was adopted from the Bader lab (https://baderlab.org/GeneSets). Similar genesets were merged together recursively to create a merged geneset if the smaller / larger geneset contains at least 50% / 80% overlapped genes. A total of 1087 genesets were applied to the model after the merging. The geneset were then normalized over genes to create the geneset-gene matrix ρ.

### Single-omic differential analyses

The gene expression level and methylation beta value readings were corrected for age and sex and normalized by mean. To balance the difference in feature number between different omics, for each omic, the inputs were normalized by multiplying a ratio of the sum of feature number of other omics over the sum of feature number of all omics.

#### Expression analysis

Differential genes included in the Bader geneset were selected by DESeq2 with P < 0.05^29^. A total of 1572 differential genes were selected.

#### Methylation analysis

Differential methylation sites proximal to at least one of the selected genes (within 500KBP) were selected with average beta value difference > 0.025 between RES and NRES. A total of 13900 differential methylation sites were selected.

#### Gene-methylation site interaction

We selected genes which were correlated with proximal methylation sites within 500KBP by Pearson correlation. For each gene, the top 10 pairs with P < 0.05 were considered as valid interaction pairs and were used to construct the interaction adjacency matrix B described in the model derivation section between gene expression and methylation.

#### Genotype analysis

Differential SNPs were identified through PLINK. A filter of Hardy-Weinberg equilibrium tests at a threshold of 0.001, missing call rate threshold at 0.01 for each SNP, and MAF threshold at 0.05 were applied, resulting in 263740 SNPs in our analysis. The SNPs were correlated with AD response and corrected by age and sex by ordinary least squares regression analysis.

### Multi-omic analyses

#### Geneset-aware NMF

To integrate the multiomic data using prior gene set information, we developed a non-negative matrix factorization (NMF) approach called geneset-aware integrative Geneset-Embedded NMF (iGEM). iGEM uses gene set information as the predefined basis matrices to help identify enriched gene sets from the gene expression and DNA methylation data. We applied iGEM to the differential genes and methylation sites selected from the single-omic differential analyse. We applied the model to derive 10 topics and the corresponding factor scores for the downstream analyses. The model design is described in the Supplementary Material.

#### Correlation analysis

Spearman correlation was performed between topic factor score W and patient condition y. Pearson correlation was performed between topic factor score W and MADRS scores. For each topic, the gene expression level and the methylation beta value from the top 10 features were correlated with MADRS score by Pearson correlation. The correlations were corrected by the Benjamini-Hochberg (BH) procedure with FDR < 0.05.

#### QTL analysis

QTL analyses were performed for the genes and methylation sites based on cis-SNPs within 500kb. In addition to gene expression and DNA methylation, QTL analysis was performed for the 10 MADRS responses and 10 topics. The QTL analyses were corrected by the BH procedure with FDR < 0.1 over in-cis SNPs.

#### Top response SNP analysis

To investigate the role of genotype variation in MDD and AD response, we have selected SNPs that were highly correlated to response with P-value < 10^−4^. Similar to the analysis previously described in the genotype analysis section, we performed regression analysis between genotype variation and patient factor score. Two sets of analyses were performed for genotype coded as either minor allele frequency or combining patients with homozygous major allele and with heterozygous patients together.

## Results

### Analysis overview

Our approach to identifying factors predicting AD treatment response is shown in **Fig. 1**. Our patient cohort consisted of 111 individuals with MDD who were treated with AD for 8 weeks, after which they were classified as responders (RES, N=61) or non-responders (NRES, N=50) based on change in Montgomery–Åsberg Depression Rating Scale (MADRS) scores between baseline and week 8. Using peripheral blood samples, we collected three high-throughput molecular datasets: RNA-sequencing, DNA methylation, and SNP genotyping. We first identified gene expression and DNA methylation differences related to response, which were used as input for our iGEM model. The model identified several sets of AD-related factors, which were investigated in downstream analyses, including their relationship to depressive symptoms, and genetic variation.

**Fig. 1.**
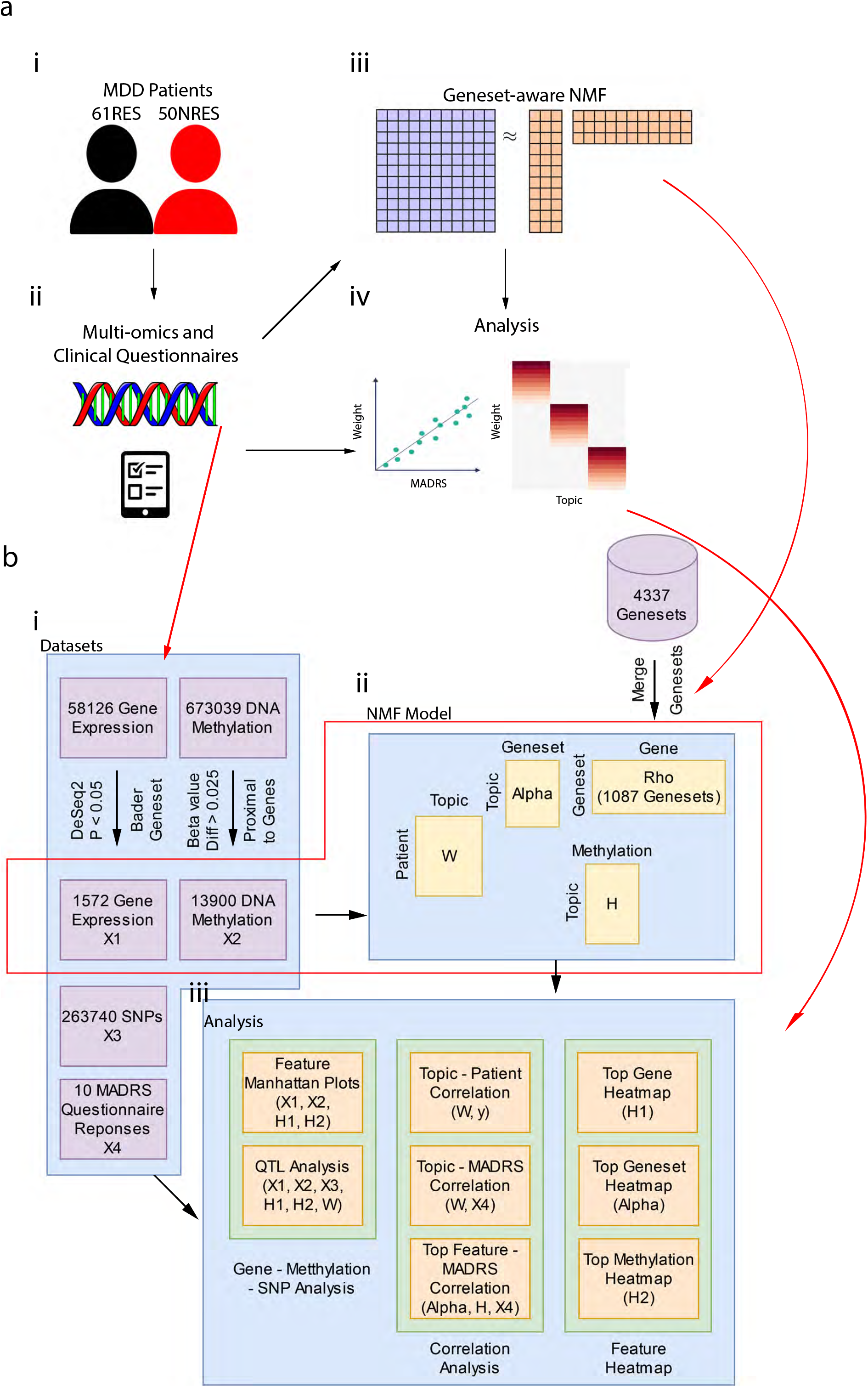
Overview of the study design. We collected blood samples and questionnaire responses from individuals with MDD who were treated for 8 weeks with SSRI or SNRIs (**Fig. 1a-i** and **Fig. 1a-ii**). The collected sample with gene expression and DNA methylation information at baseline were applied with our geneset-aware NMF (**Fig. 1a-iii**), and the derived model factor scores along with the datasets were subjected to further analysis (**Fig. 1a-iv**). In **Fig. 1b**, detailed schematics of the study design are presented and matched with the corresponding components in **Fig. 1a**. Gene expression and methylation datasets were subjected to differential analysis (**Fig. 1b-i**). The remaining differential features are applied to the model (**Fig. 1b-ii**) and the following analyses along with the model factor scores (**Fig. 1b-iii**).

### Differential analyses

We first performed differential analysis for the gene expression and DNA methylation datasets, focusing on autosomal chromosomes and protein coding genes, to identify differences related to AD response. The results are shown in **Fig. 2**. Not surprisingly, none of the features passed the stringent Bonferroni-corrected FDR of 0.05 in **Fig. 2**, partly due to the small sample size of our cohort. Nonetheless, we opted to further explore the data from a multi-omic perspective using our iGEM model by setting a lenient threshold to select as many associated candidate markers as possible. In particular, we selected genes and methylation sites that showed nominally significant differences between RES and NRES, at the p-value less than or equal to 0.05 and absolute average beta value difference greater than or equal to 0.025, for gene expression and methylation, respectively. Genesets based on the differential genes were selected from the Bader genesets described in the Methods section. A total of 1572 genes, 13900 methylation sites, 283740 SNPs, and 1087 genesets were used for our model, and downstream analyses.

**Fig. 2.**
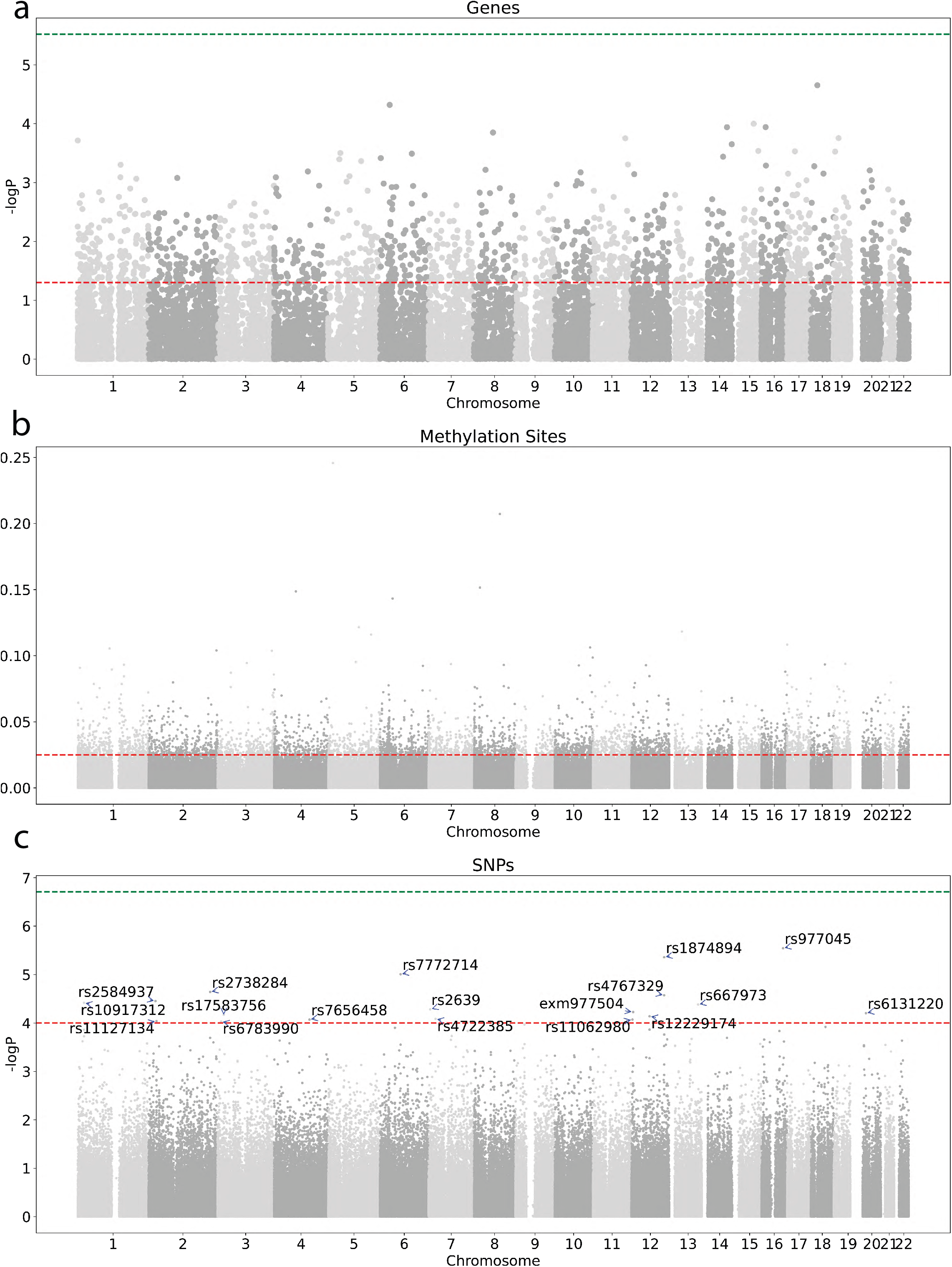
Manhattan Plots for differential analysis of gene expression (a), methylation sites (b), and SNPs (c). The green horizontal line indicates the negative log P-value where P = 0.05 / N (N = 16570, 673038, and 258782 for genes, methylation sites, and SNPs respectively). The red line indicates the differential features taken as inputs by the model (gene expression and methylation) or for further analysis (SNPs).

### Relationship of AD response with multi-omic latent factors

We applied iGEM to jointly model the filtered gene expression and methylation features. We used the Bader genesets to guide the learning of the latent factors. We performed a number of trials varying the latent factor number, and determined that 10 latent factors best captured the multi-omic dataset. We represented each subject by 10 disease factor scores, referred to as “topics”. To interpret the clinical meaning of these 10 topics, we correlated them with the AD treatment response status. We identified four significantly-correlated topics (BH-corrected FDR < 0.05). In particular, topics 5, 9, and 10 have higher factor scores in RES, while topic 4 has a higher factor score in NRES (**Fig. 3a, Supp. Table 2**). This suggests that the top gene or methylation features in topics 5,9 and 10 are associated with AD response. On the other hand, the top genes and pathways in topic 4 may be related to the AD resistance of NRES patients.

**Fig. 3.**
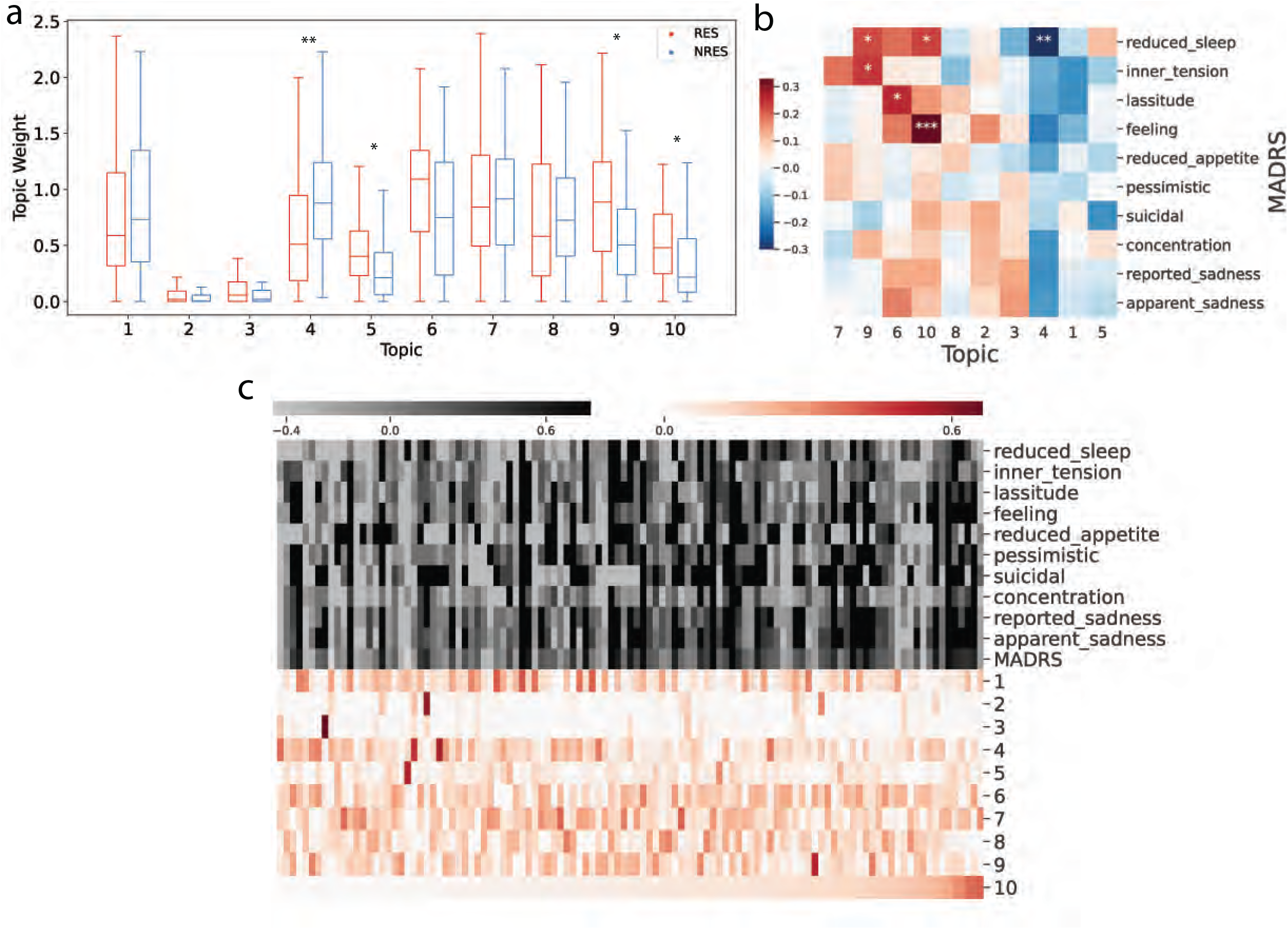
(a) Boxplot for patient factor score distribution between RES and NRES. The box indicates the range between the upper and the lower quantile. The line shows the median value. The whiskers indicate the last point within 1.5 interquantile range. *, ** represents corrected P-value < 0.05, 0.001 respectively. (b) Pearson correlation between the patient factor score and the MADRS response following treatment. The color bar shows the magnitude and sign of correlation. Topics are arranged by columns, and the MADRS responses are arranged by rows. *, **, *** represents corrected P-value < 0.01, 0.05, and 0.001 respectively. (c) Patient factor score sorted by topic 10. Each row represents a subject. The upper portion indicates how much a patient improves based on the overall MADRS score and individual MADRS response as a ratio of (week 0 score – week 8 score) / week 0 score. A greater ratio represents a greater improvement. We define RES as (week 0 score – week 8 score) / week 0 score≥ 0.5 and NRES as (week 0 score – week 8 score) / week 0 score < 0.5 respectively. A negative ratio is considered to be zero as NRES. The heatmap displays the difference between the ratio of each patient with the average value of NRES patients. The bottom portion is the patient factor score. Each column represents a topic, and the patients are sorted by the score from topic 10

We next examined the pattern of patient factor scores with respect to AD response by ordering the patients on topic 10 because due to its clinical significant and biology (described next). As shown in **Fig. 3a**, RES patients in topic 10 have a higher score than NRES patients. This is consistent with the results presented in **Fig. 3b**, where the topic 10 factor score was positively correlated with all MADRS responses.

To investigate the relationship between AD-related topics and depressive symptoms, we correlated the topics with the 10 individual MADRS items. The 10 MADRS items include “reported sadness”, “apparent sadness”, “pessimistic”, “suicidal”, “concentration”, “lassitude”, “feeling”, “inner tension”, “reduced sleep”, and “reduced appetite”. We created scores for each MADRS item by subtracting values post-AD treatment from baseline values then dividing by the baseline value, such that response to AD can be expressed by a decrease of each item, which represents a recovery from the depressed state. **Error! Reference source not found.3c** illustrates the magnitude and direction of correlation between each MADRS item and the patient factor score from the model. Five pairs were found to be significant with after correction (BH-corrected FDR < 0.05). Three topics were related to “reduced sleep”: topics 4, 9, and 10. Additionally, topic 10 was related to “feeling”, and topic 9 was related to “inner tension”. The significantly correlated pairs are shown in **Supp. Table 3**.

### Genes and genesets related to AD-response

We examined the top genes, genesets, and DNA methylation-related genes for each of the four AD-related topics (**Fig. 4**). The top geneset of topic 4 is related to the ROBO pathway, and studies have shown that upregulation of ROBO signaling is related to depression in both humans and mice^36^. On the other hand, suppression of ROBO signaling has an antidepression effect, which is mediated by an increase of neuronal cytoskeleton remodeling and neuroplasticity^37^. Topic 5 is related to integrin signaling pathways as well as several biosynthesis pathways. Topic 9 is involved with TGF-beta and AKT signaling transduction. Topic 10 is particularly interesting as it is associated with immune and inflammatory response genes and genesets. These include the top genes, *IL1RL1* and *IL5RA*, and the top genesets, *IL3* signaling pathway, *IL5* signaling pathway, and inflammatory response pathway. Indeed, previous studies have identified a relationship between AD treatment response, inflammation, and immune functioning^38, 39^. It is notable that several of the genes associated with differential methylation (*TRAPPC10, SLC1A5*, and *GPR19*) appear multiple times in the same topic, suggesting that top methylation sites are related. Additionally, *JUP* and *SLC2A5* were observed in both the top gene and top methylation sites of the same topic, which indicates a potential gene-methylation relationship within the topic.

**Fig. 4.**
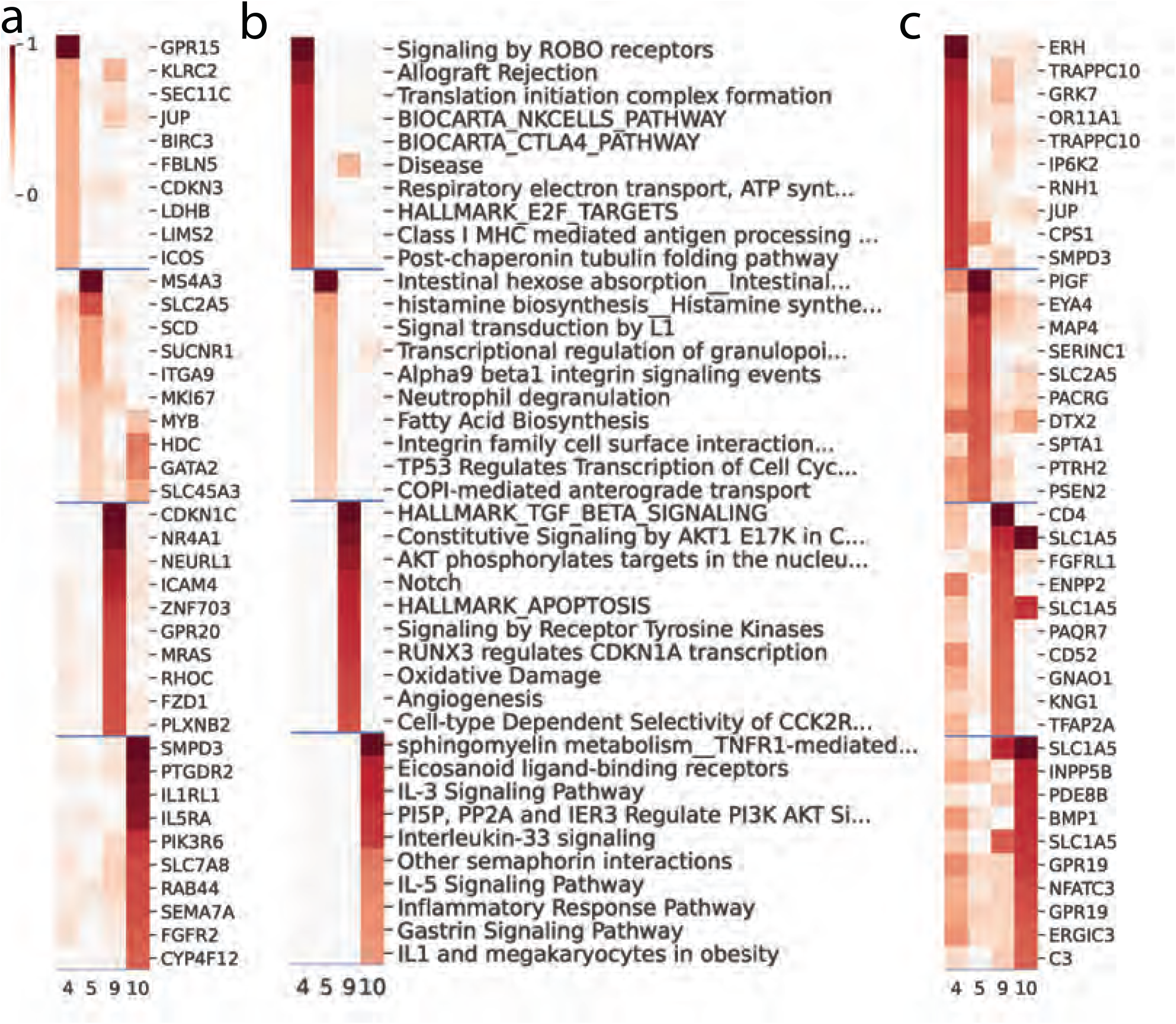
Top 10 features by feature factor score of gene (a), geneset (b), and the most correlated proximal gene of methylation sites(c). The factor scores of the features are arranged in decreasing order of rank from top to bottom. The top 10 rows are from the top 10 features in the first topic. The next 10 rows are the top features from the next topic, and so on. For methylation, the names of the top methylation sites were substituted with its most correlated gene among proximal genes within 500kb, determined by Pearson correlation. The color represents the relative score of a feature in each topic within a range of 0-1. The top feature in each topic has a score of 1, and the score of other features are presented as a proportion of the top feature score within the same topic.

### Association of top genes under AD-related topics with specific depressive symptoms

As the diagnosis of MDD requires the presence of any five diagnostic criteria out of a list of symptoms, provided that one of the criteria is presence of depressed mood or lack of interest, the symptoms used to diagnose depression may vary from individual to individual, we set out to determine if there were any molecular markers that associate with a given symptom. For each topic, we correlated the top 10 genes with each individual MADRS item. The significant pairs after correction for multiple testing (BH-corrected FDR < 0.1) are listed in **Error! Reference source not found**.. Among the 10 MADRS responses, we found “reduced sleep”, “inner tension”, and “feeling” to be significantly associated with at least one gene. Three genes under topic 10 were significantly correlated with two MADRS responses at FDR < 0.05: RAB44 -reduced sleep, SEMA7A - feeling, and IL5RA - feeling. At the more lenient threshold of FDR < 0.1, we found additional associations, including (in the format of topic-MADRS-gene(s)): topic 4 - reduced sleep - BIRC3/LIMS2/LDHB/ICOS, topic 5 - feeling - GATA2/SLC45A3, topic 5 - reduced sleep - GATA2/HDC, topic 9 - reduced sleep - NR4A1/NEURL1, topic 10 - reduced sleep - SMPD3/PTGDR2/PIK3R6/SEMA7A/IL1RL1/SLC7A8/CYP4F12, and topic 10 - feeling - CYP4F12/PTGDR2/PI3KR6/SMPD3/RAB44.

### Multi-omic quantitative trait loci analysis

To investigate the impact of genotype variation on AD response, we performed cis-eQTL and cis-mQTL analyses for the differential genes and methylation sites. We tested the top 10 genes under the AD-related topics and discovered several eQTL for genes *SLC45A3, ICAM4, PLXNB2*, and *PIK3R6*, which are among the top genes under topics 5, 9, 9, and 10, respectively (**Supp. Table 5**). Interestingly, *PLXNB2* each harbor an eQTL SNP in their regions. Using the top methylation sites under each topic, we also identified a substantial number of mQTL (**Supp. Table 6**), implying the epigenetic associations with AD-response.

The results are presented in Manhattan plots, where genes and methylation sites are aligned by genomic location to display the in-cis relationships between the top features identified by the model (**Supp. Figure 4**). We found several top methylation sites that were proximal to one of the top genes with P-value < 0.05: Topic 4 - JUP - cg11278727, topic 5 - SLC2A5 - cg04413644, topic 5 - SLC2A5 - cg25755851, topic 5 - MYB - cg04531756, topic 10 - IL1RL1 - cg03938978, and topic 10 - CYP4F12 - cg24656834.

Finally, we performed regression analysis between the 18 top response SNPs, shown in **Fig. 2C** (P-value < 10^−4^) and AD response-associated topics. The t-statistics of the regression analyses are shown as heatmaps in **Supp. Figure 5**. Not all SNPs are significantly correlated with the differential topics, and interestingly, each topic displayed a unique pattern of association with these SNPs. We identified 7 significant SNP-topic pairs, as shown in **Table 1**. Interestingly, several of these SNPs are located near genes which have previously been associated with MDD. Finally, in order to better characterize the effects of the minor allele, we grouped individuals who were heterozygous with those who were homozygous at the major allele and repeated this analysis. These results are shown in (**Supp. Table 7**).

**Table 1.**
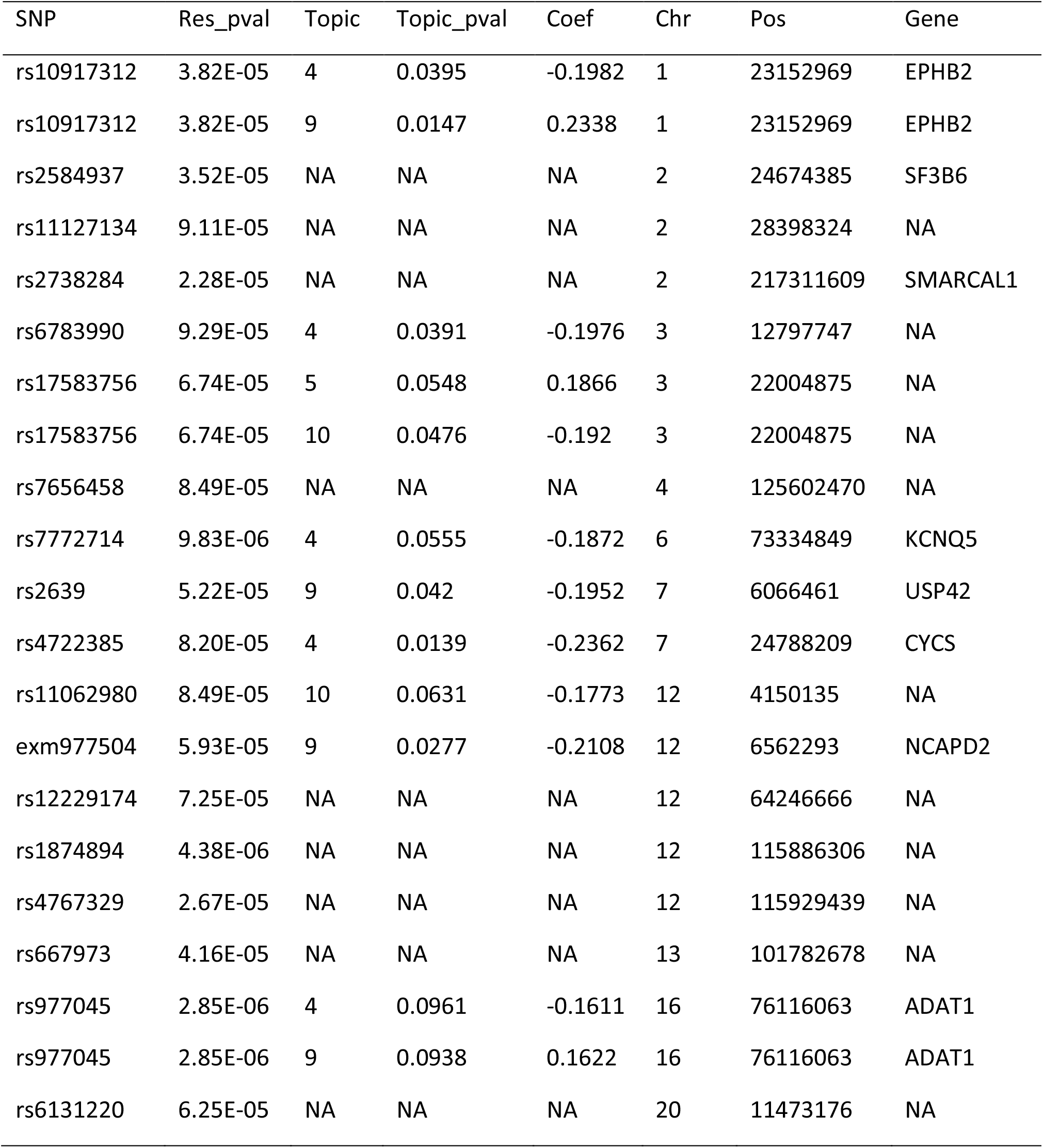
Relationship between SNPs associated with response and response-associated topics. The associations between top SNPs and the response are indicated by the P-value (Res_pval). In cases where differential topics are associated with the SNPs (P-value < 0.1), P-value (Topic_pval) and correlation coefficient (Coef) are indicated. The chromosome (Chr) and genomic location (Pos) of each SNP are shown. The closest differentially expressed genes within 500kb are shown.

## Discussion

Currently, the molecular factors underlying response to AD treatment remain unknown, which impacts our capacity to better develop personalized treatment strategies for depressed patients, as well as to develop new treatment avenues. While early studies investigated candidate genes or small panels of potential biomarkers, more recent studies have investigated genome-wide data, with the expectation that the combination of numerous biological measures may have a better capacity to predict or explain response. In the present study, we combined several genome-wide datasets in order to identify underlying biological characteristics related to AD response, as well as to identify biological pathways which may be relevant. Using the iGEM model, we identified four topics that were significantly associated with response. These topics comprised genes displaying differential expression between responders and non-responders, genes associated with DNA methylation differences, and genesets associated with these top genes. Interestingly, the topic that was most positively correlated with AD response was associated with immune and inflammatory functioning, both of which have been associated with MDD.

Several of the top genes identified using the model to examine gene expression and methylation data are related to depressive symptoms, and have previously been found to be differentially expressed in the context of depression or associated with AD response. In particular, we have identified IL-related genes interleukin 1 receptor like 1 (IL1RL1) and interleukin 5 receptor (IL5) subunit alpha (IL5RA) as well as the AKT/PIK3 pathway related gene phosphoinositide-3-kinase regulatory subunit 6 (PIK3R6). A previous study found that IL5RA and PIK3R6 were upregulated in lithium-treated bipolar disorder patients^40^. IL5 signaling affects the downstream AKT/PIK3 pathway, which suggests that AD response could be mediated in a top-down fashion from IL5 to the AKT/PIK3 pathway and other downstream molecules^41^. IL1RL1 mRNA was found to be differentially expressed in the spleen of mice with depression-like behaviour^42^. In addition, the expression of IL1RL1 (also known as ST2 and IL33R), has been shown to be negatively associated with depression severity in post-stroke depression, whereas the expression of its ligand, the cytokine IL33, was positively associated with severity^43-46^. In addition to the inflammatory regulators, another top gene in topic 10, sphingomyelin phosphodiesterase 3 (SMPD3), has also been associated with depression and AD treatment response. Acid sphingomyelinase (ASM) is a molecular target of AD drugs, and mice treated with the SSRI fluoxetine demonstrated inhibited ASM activity^47^. ASM could potentially mediate its effect through ceramide and sphingolipid-metabolizing enzymes including SMPD3, and ASM overexpression led to reduced SMPD3 expression in mice, which was accompanied by an increase of depression-like behaviour^48, 49^.

We included genotype information as a secondary set of analyses, and identified a number of SNPs which were related to response, including several which were also associated with the AD-related topics. Interestingly, several of the genes proximal to these SNPs have previously been associated with depression, including nuclear receptor coactivator 1 (NCOA1), which is 40398 bp downstream of rs2584937^50^. NCOA1 is involved in glucocorticoid receptor signaling, and impairments of glucocorticoid receptor signaling have previously been associated with MDD^51^. Moreover, the increase of NCOA1 during pregnancy was diminished in depressed patients, which also suggests an association between depression and NCOA1^52^. Also of note is the relationship between response and exm977504 (rs2041385), which is found within TAP Binding Protein Like (TAPBPL), an MHC-related gene which has previously been shown to be influenced by glucocorticoid signaling^53^. This SNP is an eQTL for TAPBPL expression in various tissues including blood, brain, LCL, and monocytes^50, 54-57^. Interestingly, additional eQTLs were found for TAPBL in an MDD population, however this association was only significant in females^53^. We also identified SNPs which were near genes which were differentially expressed in relation to AD response, and which have previously been associated with MDD. rs10917312 is near a member of receptor tyrosine kinase transmembrane glycoproteins, EPH Receptor B2 (EPHB2). KO mice with EPHB2 deficiency have display depressive and stressed behaviours^58, 59^. Additionally, the deficiency in human APP/PS1 transgenic mice can be alleviated with overexpression of EPHB2 with improved impaired memory, depression, and anxiety-like behaviours^60^. Another gene to be noted is Potassium Voltage-Gated Channel Subfamily Q Member 5 (KCNQ5), which is located near rs7772714. KCNQ5 was downregulated along with several other genes in the cholinergic synapse pathway in individuals with MDD^61^. Furthermore, the expression of KCNQ5 expression was upregulated in mice treated with amitriptyline, which indicates a potential role of signaling transduction in the AD response^62^. Although we did not identify in-cis associations between the top response SNPs and the top features in differential topics, the topic-SNP-gene relationships identified from our analyses complement our findings with iGEM, and shed light on the complex relation between various genes associated with MDD pathogenesis and AD response.

The proposed iGEM model has several advantages. As previously mentioned, AD response in MDD is a highly complex biological process, that likely involves, and is influenced by, numerous genetic and epigenetic processes. One of the advantages of the model is that it promotes the finding of multiple features with similar functions by integrating geneset information. In addition to gene expression and DNA methylation, the model is flexible and may accept different types of biological readings. A potential usage may include metabolite readings. If there are known associations between metabolites and a particular disease, these can be included in a similar fashion as the geneset information in the current model. There are several limitations of the model that must be noted. Currently, the model only supports two types of data input, but has the potential to extend to three or more types if needed. Another limitation of the model is that, as an NMF-based model, it takes only non-negative inputs. Therefore, it may be difficult to apply the model to certain data types that contain negative values, which requires proper normalization before applying the model.

In summary, we found biologically meaningful features related to MDD and AD response, which may ultimately lead to not only the ability to predict AD response, but also the development of new treatments for MDD. With the auxiliary geneset information, the model promotes findings of biological relevance and intuitive interpretation of latent topic based on the associated genesets. For example, the immune and inflammatory processes highlighted by topic 10, as well as the association of several response-associated SNPs with genes and pathways which have previously been associated with MDD.

Overall, the model identified several biological differences between individuals who responded to AD treatment and those who did not, and emphasizes the involvement of immune system in AD treatment response. Future studies are needed to better characterize the role of these molecular mechanisms in AD response, as well as to determine how well these findings extend to other cohorts.

## Acknowledgements

This work was supported by the Canadian Institutes of Health Research (CIHR) (grant no. FRN/MOP#111260) as well as Janssen Research & Development. Y.L. is supported by Natural Sciences and Engineering Research Council (NSERC) Discovery Grant (RGPIN-2019-0621), Fonds de recherche Nature et technologies (FRQNT) New Career (NC-268592), and Canada First Research Excellence Fund Healthy Brains for Healthy Life (HBHL) initiative New Investigator start-up award (G249591).

## Supplementary Information

Supplementary information is available at the Molecular Psychiatry website.

## Figures

**Supp. Figure 1.**
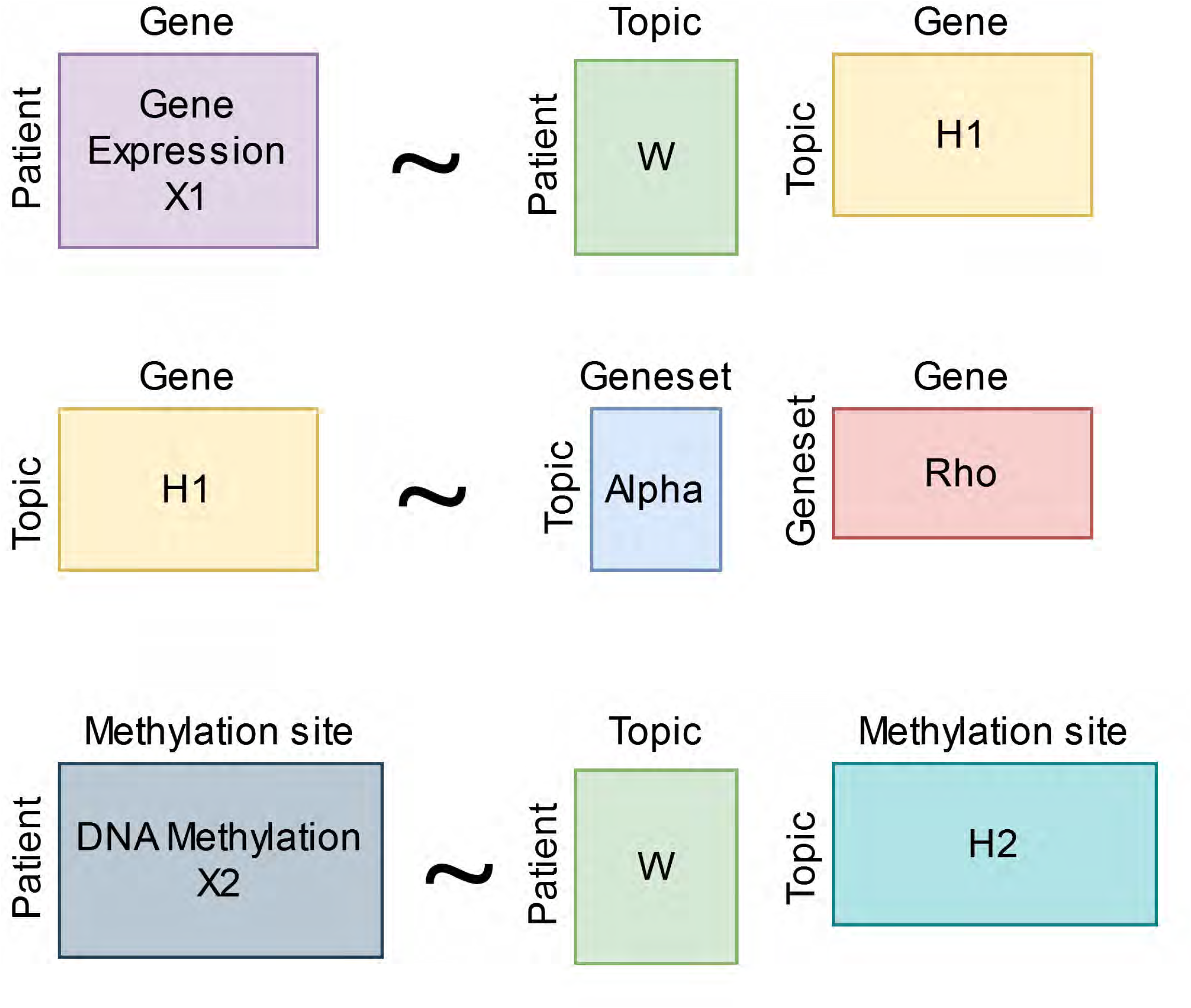
Overview of the model design. Matrix decomposition in the model can be separated into the 3 steps illustrated in the figure.

**Supp. Figure 2.**
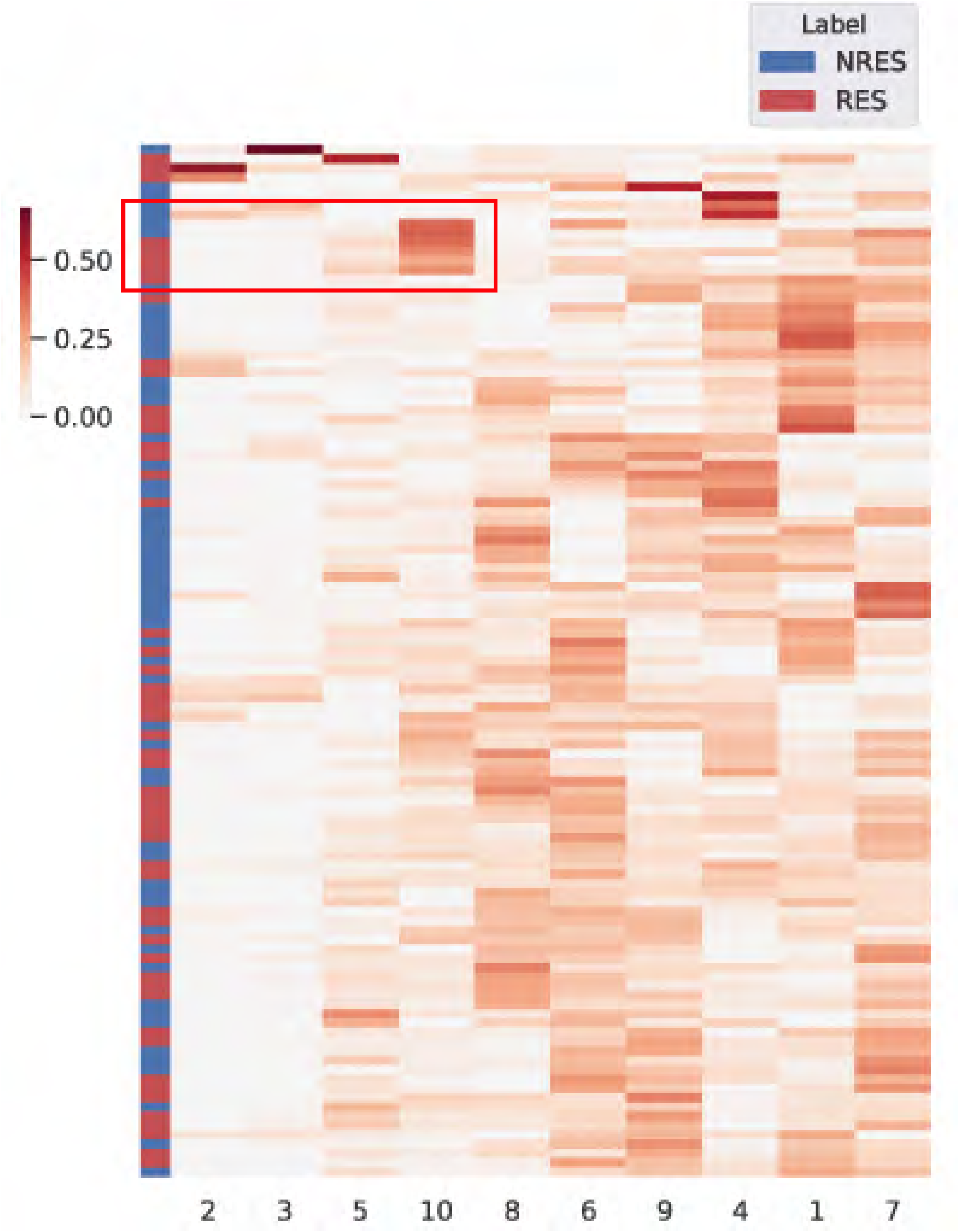
Hierarchical clustering of patient factor score. Each row represents a subject. The left portion indicates whether a patient is RES (red) or NRES (blue) based on the overall MADRS score. The right portion is the patient factor score. Each column represents a topic. The rows and columns are sorted by hierarchical clustering, with similar topics and subjects arranged closely. For differential topics, certain pattern of patient clustering can be observed. The red box indicates a patient cluster with higher topic 10 score.

**Supp. Figure 3.**
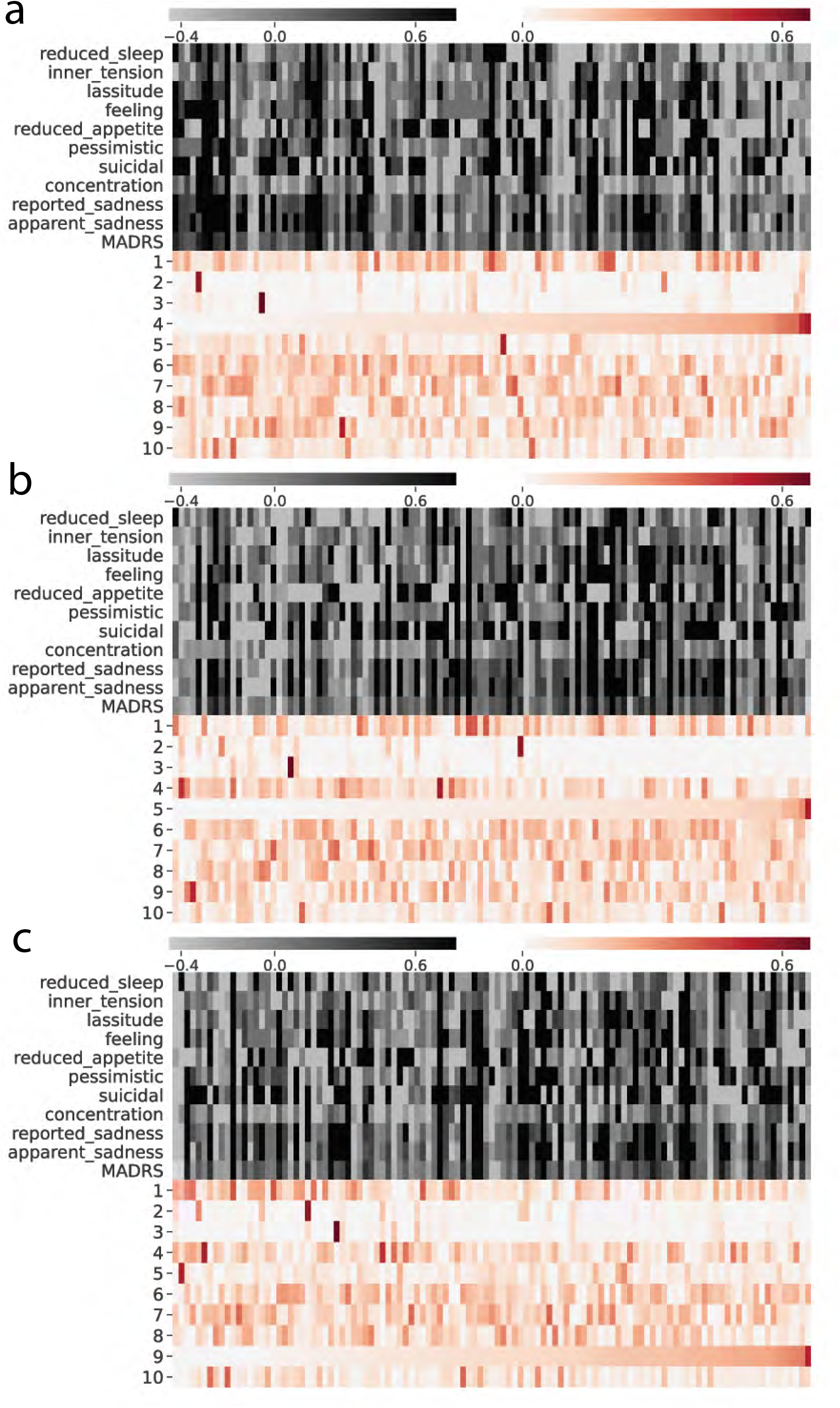
Patient factor score sorted by topic 4(a), 5(b), and 9(c).

**Supp. Figure 4.**
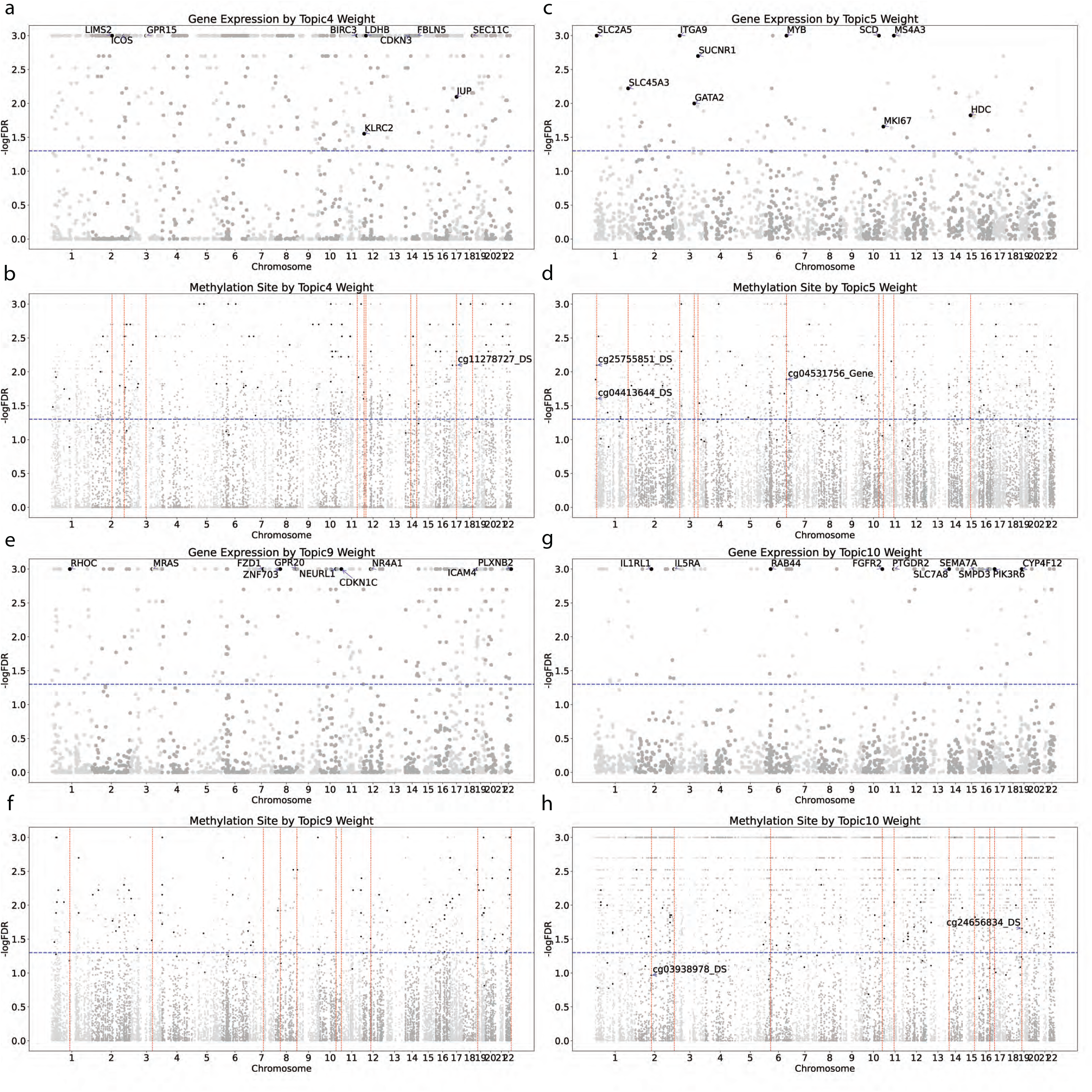
Gene and methylation Manhattan plots. The Manhattan plots for gene expression and methylation of topic 4 (a. and b.), 5 (c. and d.), 9 (e. and f.), and 10 (g. and h.). Feature Manhattan plots are generated by permutation tests on feature factor scores with data shuffled over patients for 1000 models with fixed model factor score and identical hyperparameters based on the tested model. For each topic, the patient factor scores were tested against SNP variation by an OLS regression model, and the P-values were corrected by the BH procedure. Features are arranged by genomic location. Y axis shows the negative log P-value after permutation test. The blue horizontal line indicates the negative log P-value where P = 0.05. The red vertical line indicates the genomic location of top genes. In the top panel, the top 10 genes are highlighted in black with the name labeled. In the bottom panel, the top 100 methylation sites are highlighted in black. Among the top 100 sites, the ones proximal to the top genes have the name labeled with the relative position to the top genes (US, DS, and Gene for upstream, downstream, and within gene transcript respectively).

**Supp. Figure 5.**
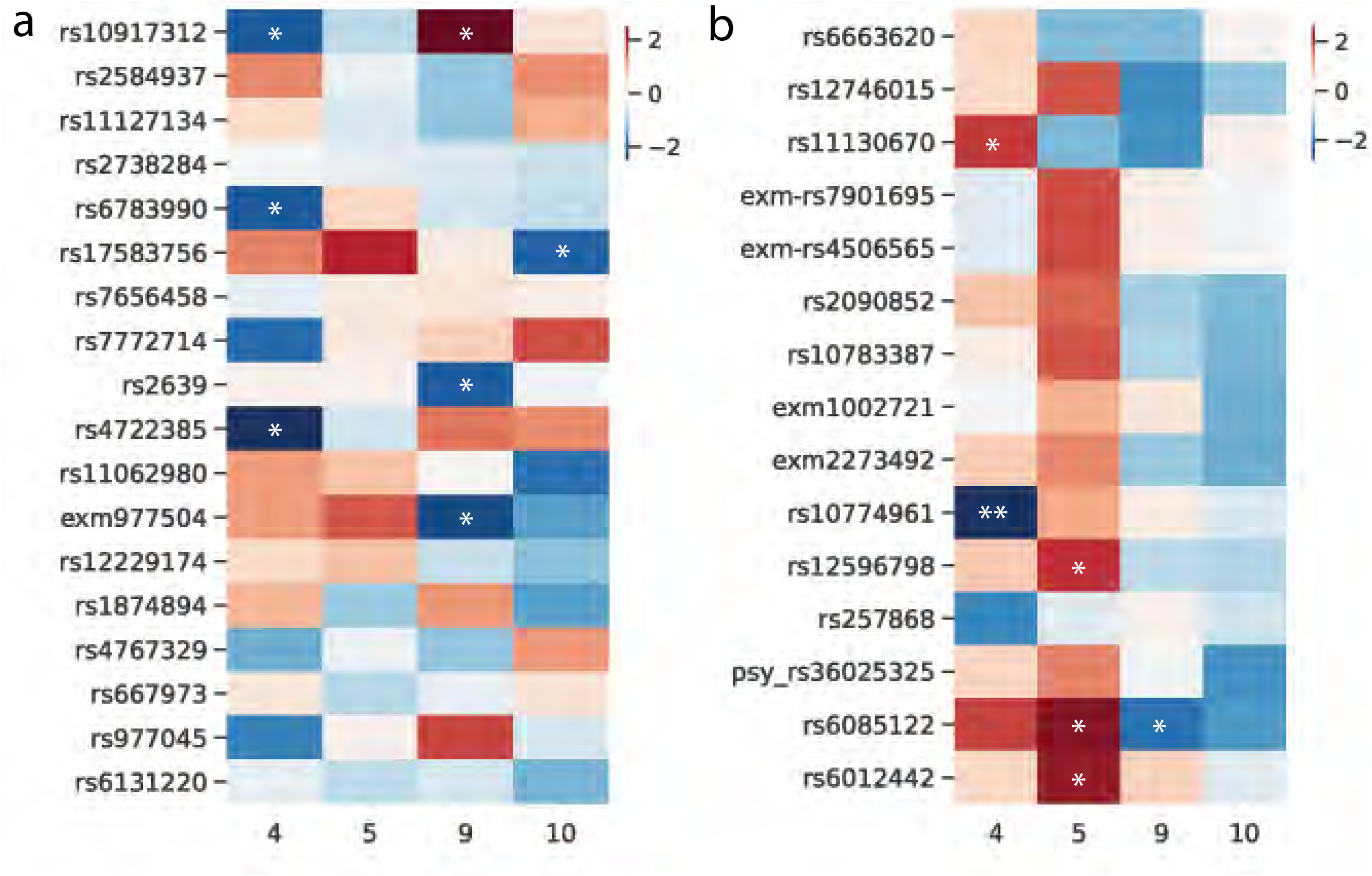
Heatmap of t-statistics for correlation between top response SNPs and differential topics with minor allele frequency (a.) and alternative genotype grouping (b.) as described in **Methods**. The asterisks * and ** indicate P-value < 0.05 and < 0.01 respectively.

## Supplementary materials

### Integrative Geneset-Embedded non-negative Matrix factorization (iGEM)

#### Model Loss

With the squared error loss function L, a number of k multi-omic datasets can be denoted as X (patient by feature) and reconstructed as matrices patient factor score W (patient by topic) and feature factor score H (topic by feature), given n features, m topics, and p patients. e is a vector of ones / a matrix with ones on the diagonal and zeros elsewhere. The sparsity constraint is performed as the square of L1-norm on the column vector h_j_, which is the j-th column of H. Here, we denote the squared error loss and the sparsity constraint as L_a_.

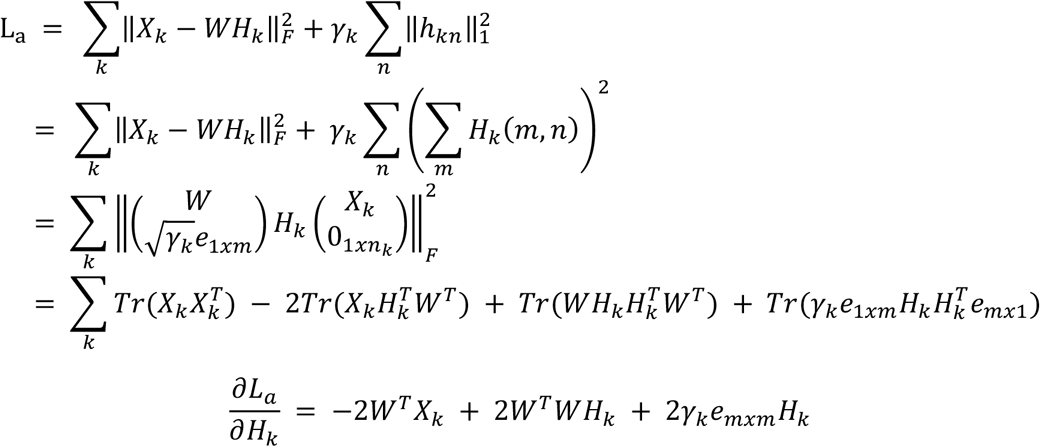

In addition to the sparsity constraint, a rewarding term can be added to strengthen the entries involving the interaction within each dataset or between different datasets. The rewarding terms, with the strength determined by the coefficients λ_k_ and λ_kj_, are added as a trace between the feature factor scores H and the adjacency matrix. The adjacency matrices A and B indicate the interactions within α_k_ and between α_k_ and another dataset H_j_ respectively. For the patient factor score W, a penalty term 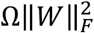is added to limit its increase. We combine these constraints as L_b_.

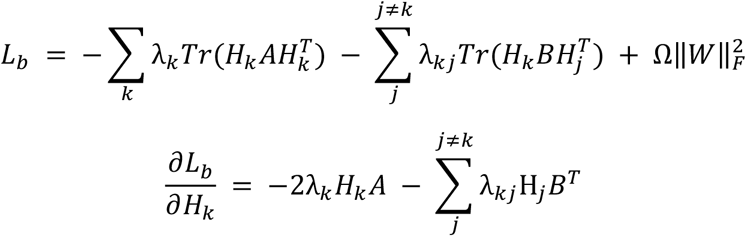

For a dataset such as gene expression with additional information of gene composition in genesets, it is possible to derive gene and geneset information with shared topic. The matrix H (topic by gene) can be reconstructed by matrices α (topic by geneset) and ρ (geneset by gene), with a number of l genesets. The squared error loss function L_c_ with sparsity constraint on α can be written in a similar fashion as L_a_.

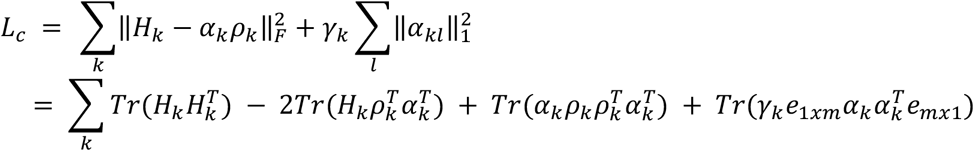

The loss function of the model L is composed of L_a_, L_b_, and L_c_:

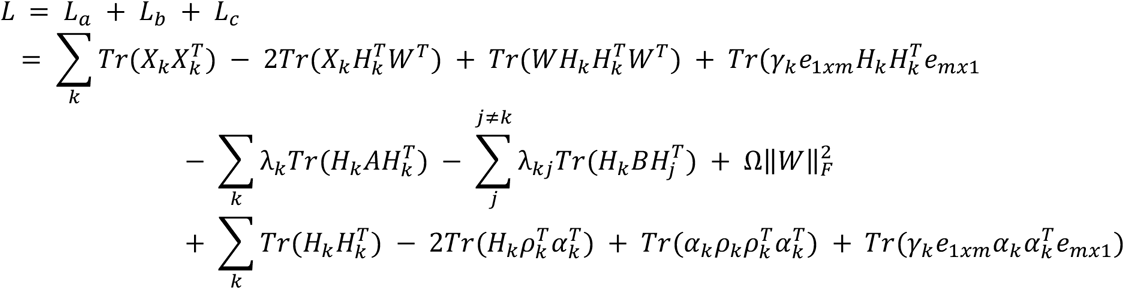

#### Multiplicative Update

For the patient factor score W, the partial derivative of L with respective to W can be written as 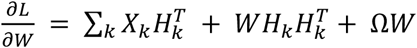. Note that a common numerical constant of 2 are removed for brevity, and the same will be applied to the rest of the partial derivative steps. The multiplicative update of W can be performed with a learning rate 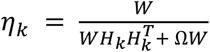.

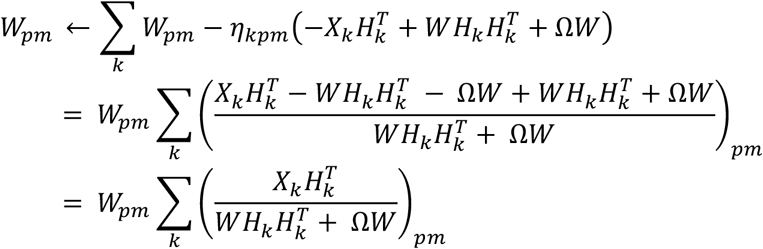

For a geneset factor score α_k_, the partial derivative of L with respective to α_k_ can be written as 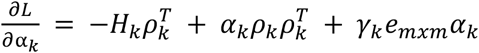. The multiplicative update of α_k_ can be performed with a learning rate 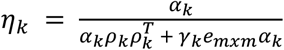.

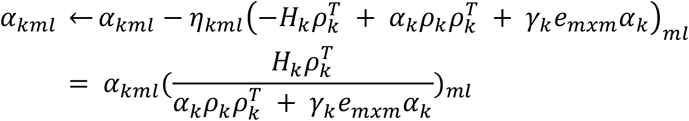

For a feature factor score α_k_, the partial derivative of L with respective to α_k_ can be written as 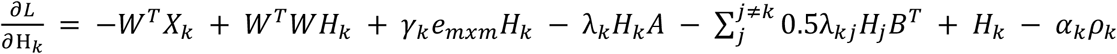. The multiplicative update of α_k_ can be performed with a learning rate 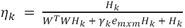.

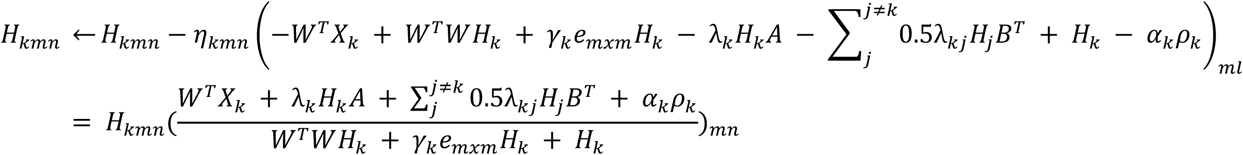

#### Model update steps

1. Initialize W, H, and α.

2. Fix H and update W with 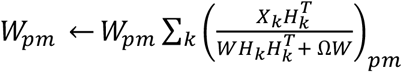
(Solve 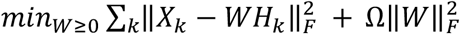)

3.Fix W, α and update H with 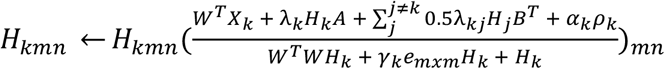
(Solve 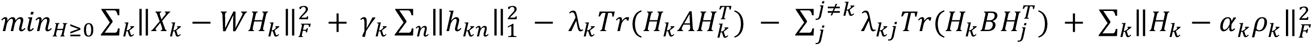)

4. Fix H and update α with 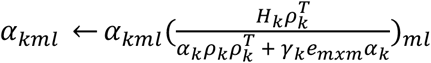
(Solve 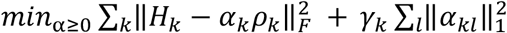)

5. Repeat 2-4 until convergence or reaching maximum number of iterations.

#### Model settings

There are several hyperparameters that affect the model performance. The reward term between omics λ_kj_ is set at 0.01. The penalty term Ω for patient factor score W is set at 2E^-6^. The sparsity constraint γ for gene, geneset, and methylation are set at 0.1, 0.05, and 0.01 respectively.

**Supp. Table 1.** Demographic characteristics of cohort. Individuals with major depressive disorder were classified as treatment non-responders (NRES) or responders (RES) based on a ≤ 50% reduction in total Montgomery-Åsberg Depression Rating Scale (MADRS) score between baseline (T0) and after 8 weeks of treatment (T8) with either escitalopram or desvenlafaxine. Values are reported as average ± standard error.

|  | **NRES** | **RES** |
| --- | --- | --- |
| N | 50 | 61 |
| Age | 41.0 ± 1.8 | 38.6 ± 1.6 |
| Sex (% female) | 60.0% | 65.6% |
| Medication (% escitalopram) | 56.0% | 44.3% |
| MADRS (T0) | 34.2 ± 0.8 | 31.9 ± 0.8 |
| MADRS (T8) | 26.5 ± 1.1 | 8.9 ± 0.7 |

**Supp. Table 2.** Spearman correlation between the patient factor score and the patient condition. Each row represents the Spearman correlation coefficient (Coef.), P-value, and corrected P-value after multiple testing (BH) of a topic.

| Topic | Coef. | P-value | BH |
| --- | --- | --- | --- |
| 4 | 0.3165 | 0.0007 | 0.0071 |
| 10 | -0.2729 | 0.0038 | 0.0188 |
| 9 | -0.2577 | 0.0063 | 0.0211 |
| 5 | -0.2283 | 0.016 | 0.0399 |
| 6 | -0.1916 | 0.044 | 0.088 |
| 2 | -0.1503 | 0.1153 | 0.1922 |
| 3 | -0.1368 | 0.1524 | 0.2177 |
| 1 | 0.1051 | 0.2722 | 0.3403 |
| 8 | 0.022 | 0.8184 | 0.9093 |
| 7 | 0.0085 | 0.9296 | 0.9296 |

**Supp. Table 3.** Pearson correlation between the patient factor score and the MADRS response. Each row represents the Pearson correlation coefficient (Coef.), P-value, and corrected P-value after multiple testing between a MADRS response and a topic.

| MADRS | Coef. | P-value | BH | Topic |
| --- | --- | --- | --- | --- |
| feeling | 0.3571 | 0.0001 | 0.0012 | 10 |
| reduced_sleep | 0.3335 | 0.0003 | 0.0017 | 10 |
| reduced_sleep | -0.2531 | 0.0074 | 0.0736 | 4 |
| inner_tension | 0.2469 | 0.009 | 0.0899 | 9 |
| lassitude | 0.2242 | 0.018 | 0.1799 | 6 |
| reported_sadness | 0.2195 | 0.0206 | 0.1091 | 5 |
| apparent_sadness | 0.2175 | 0.0218 | 0.1091 | 5 |
| apparent_sadness | -0.202 | 0.0335 | 0.1001 | 4 |
| feeling | -0.2007 | 0.0347 | 0.1001 | 4 |
| reported_sadness | -0.1952 | 0.0401 | 0.1001 | 4 |

**Supp. Table 4.**
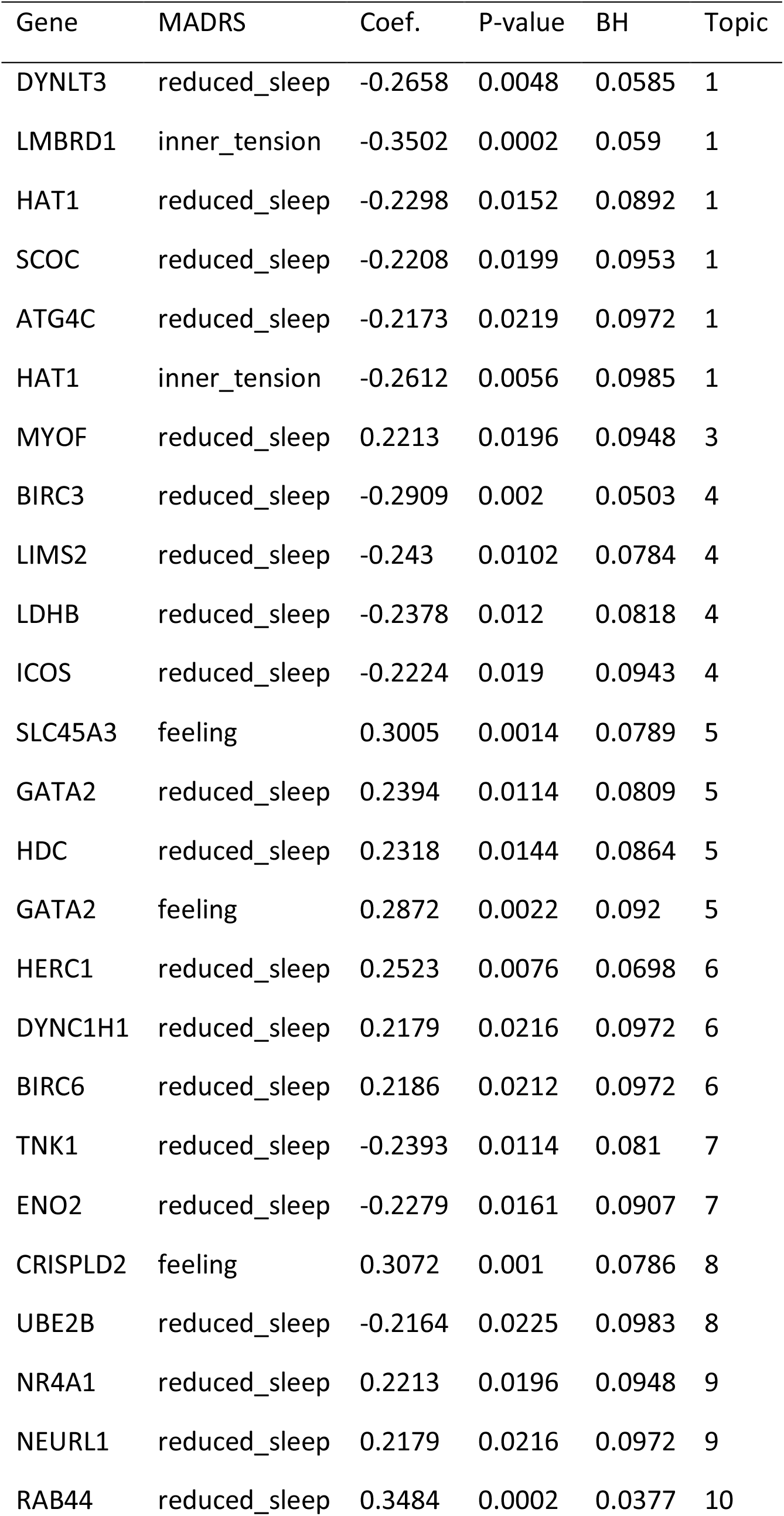
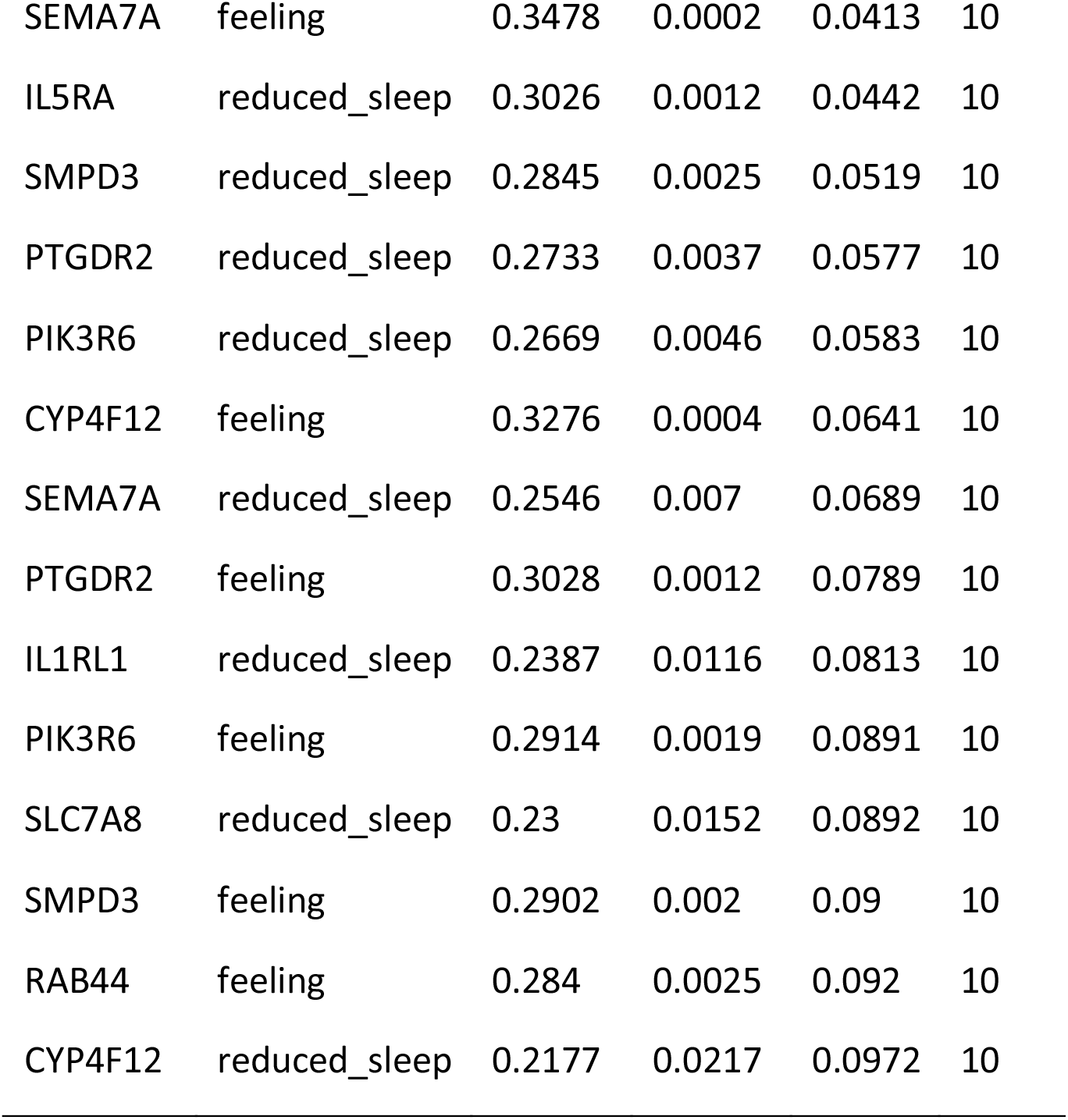
Pearson Correlation between the gene factor score and the MADRS response. Each row represents the Pearson correlation coefficient (Coef.), P-value, and corrected P-value after multiple testing between a top gene from the corresponding topic and MADRS response.

| Gene | MADRS | Coef. | P-value | BH | Topic |
| --- | --- | --- | --- | --- | --- |
| DYNLT3 | reduced_sleep | -0.2658 | 0.0048 | 0.0585 | 1 |
| LMBRD1 | inner_tension | -0.3502 | 0.0002 | 0.059 | 1 |
| HAT1 | reduced_sleep | -0.2298 | 0.0152 | 0.0892 | 1 |
| SCOC | reduced_sleep | -0.2208 | 0.0199 | 0.0953 | 1 |
| ATG4C | reduced_sleep | -0.2173 | 0.0219 | 0.0972 | 1 |
| HAT1 | inner_tension | -0.2612 | 0.0056 | 0.0985 | 1 |
| MYOF | reduced_sleep | 0.2213 | 0.0196 | 0.0948 | 3 |
| BIRC3 | reduced_sleep | -0.2909 | 0.002 | 0.0503 | 4 |
| LIMS2 | reduced_sleep | -0.243 | 0.0102 | 0.0784 | 4 |
| LDHB | reduced_sleep | -0.2378 | 0.012 | 0.0818 | 4 |
| ICOS | reduced_sleep | -0.2224 | 0.019 | 0.0943 | 4 |
| SLC45A3 | feeling | 0.3005 | 0.0014 | 0.0789 | 5 |
| GATA2 | reduced_sleep | 0.2394 | 0.0114 | 0.0809 | 5 |
| HDC | reduced_sleep | 0.2318 | 0.0144 | 0.0864 | 5 |
| GATA2 | feeling | 0.2872 | 0.0022 | 0.092 | 5 |
| HERC1 | reduced_sleep | 0.2523 | 0.0076 | 0.0698 | 6 |
| DYNC1H1 | reduced_sleep | 0.2179 | 0.0216 | 0.0972 | 6 |
| BIRC6 | reduced_sleep | 0.2186 | 0.0212 | 0.0972 | 6 |
| TNK1 | reduced_sleep | -0.2393 | 0.0114 | 0.081 | 7 |
| ENO2 | reduced_sleep | -0.2279 | 0.0161 | 0.0907 | 7 |
| CRISPLD2 | feeling | 0.3072 | 0.001 | 0.0786 | 8 |
| UBE2B | reduced_sleep | -0.2164 | 0.0225 | 0.0983 | 8 |
| NR4A1 | reduced_sleep | 0.2213 | 0.0196 | 0.0948 | 9 |
| NEURL1 | reduced_sleep | 0.2179 | 0.0216 | 0.0972 | 9 |
| RAB44 | reduced_sleep | 0.3484 | 0.0002 | 0.0377 | 10 |
| SEMA7A | feeling | 0.3478 | 0.0002 | 0.0413 | 10 |
| IL5RA | reduced_sleep | 0.3026 | 0.0012 | 0.0442 | 10 |
| SMPD3 | reduced_sleep | 0.2845 | 0.0025 | 0.0519 | 10 |
| PTGDR2 | reduced_sleep | 0.2733 | 0.0037 | 0.0577 | 10 |
| PIK3R6 | reduced_sleep | 0.2669 | 0.0046 | 0.0583 | 10 |
| CYP4F12 | feeling | 0.3276 | 0.0004 | 0.0641 | 10 |
| SEMA7A | reduced_sleep | 0.2546 | 0.007 | 0.0689 | 10 |
| PTGDR2 | feeling | 0.3028 | 0.0012 | 0.0789 | 10 |
| IL1RL1 | reduced_sleep | 0.2387 | 0.0116 | 0.0813 | 10 |
| PIK3R6 | feeling | 0.2914 | 0.0019 | 0.0891 | 10 |
| SLC7A8 | reduced_sleep | 0.23 | 0.0152 | 0.0892 | 10 |
| SMPD3 | feeling | 0.2902 | 0.002 | 0.09 | 10 |
| RAB44 | feeling | 0.284 | 0.0025 | 0.092 | 10 |
| CYP4F12 | reduced_sleep | 0.2177 | 0.0217 | 0.0972 | 10 |

**Supp. Table 5.** eQTL Result of top 10 genes in the differential topics. The columns include the topic, the chromosome (Chr), the gene name and its genomic location (Gene, Gene_Start, Gene_End), the SNP name and its genomic location (SNP, SNP_Loc), QTL correlation statistics including P-Value, correlation coefficient, and BH-corrected FDR (P-value, Coef, BH) were included.

| Topic | Chr | Gene | Gene_Start | Gene_End | SNP | SNP_Loc | P-value | Coef | BH |
| --- | --- | --- | --- | --- | --- | --- | --- | --- | --- |
| 5 | 1 | SLC45A3 | 205626979 | 205649637 | rs913722 | 205318983 | <0.0001 | -0.3115 | 0.0042 |
| 9 | 19 | ICAM4 | 10397631 | 10399198 | rs875569 | 10668953 | 0.0026 | 0.2048 | 0.0928 |
| 9 | 19 | ICAM4 | 10397631 | 10399198 | exm1426306 | 10670992 | 0.0021 | 0.2115 | 0.0928 |
| 9 | 19 | ICAM4 | 10397631 | 10399198 | rs12984043 | 10672493 | 0.0003 | 0.2609 | 0.0331 |
| 9 | 22 | PLXNB2 | 50713408 | 50746075 | exm2235942 | 50713409 | 0.0011 | -0.2123 | 0.0866 |
| 9 | 22 | PLXNB2 | 50713408 | 50746075 | exm1621884 | 50716167 | 0.0002 | -0.2403 | 0.0283 |
| 10 | 17 | PIK3R6 | 8706040 | 8770994 | rs6503145 | 8613462 | 0.0019 | 0.2148 | 0.0848 |
| 10 | 17 | PIK3R6 | 8706040 | 8770994 | rs2101938 | 8620629 | 0.0015 | -0.2386 | 0.0848 |
| 10 | 17 | PIK3R6 | 8706040 | 8770994 | rs8071193 | 8655415 | 0.0015 | 0.2209 | 0.0848 |

**Supp. Table 6.** mQTL Result of top 10 methylation sites in the differential topics. The columns include the topic, the chromosome (Chr), the methylation site name and its genomic location (Met, Met_Start), the SNP name and its genomic location (SNP, SNP_Loc), QTL correlation statistics including P-Value, correlation coefficient, and BH-corrected FDR (P-value, Coef, BH) were included. The closest proximal gene to the methylation site and SNP were also included (Met_Gene and SNP_Gene)

| Topic | Chr | Met | Met_Start | SNP | SNP_Loc | Met_Gene | SNP_Gene | P-value | Coef | BH |
| --- | --- | --- | --- | --- | --- | --- | --- | --- | --- | --- |
| 4 | 3 | cg05107650 | 141441554 | rs1026716 | 140983891 | GRK7 | None | 0.0051 | 0.0521 | 0.0875 |
| 4 | 3 | cg05107650 | 141441554 | psy_rs12491228 | 141031422 | GRK7 | GRK7 | 0.0069 | 0.0593 | 0.0989 |
| 4 | 3 | cg05107650 | 141441554 | rs9825379 | 141137035 | GRK7 | GRK7 | 0.0004 | 0.1089 | 0.0189 |
| 4 | 3 | cg05107650 | 141441554 | exm-rs10513137 | 141143430 | GRK7 | GRK7 | 0.0023 | 0.088 | 0.0659 |
| 4 | 3 | cg05107650 | 141441554 | rs6762826 | 141145315 | GRK7 | GRK7 | 0.0045 | 0.0783 | 0.0875 |
| 4 | 3 | cg05107650 | 141441554 | rs295322 | 141326602 | GRK7 | GRK7 | 0.0001 | 0.1144 | 0.0046 |
| 4 | 16 | cg09821790 | 68283575 | psy_rs77517043 | 68301410 | NFATC3 | NFATC3 | 0.0069 | 0.0319 | 0.05 |
| 4 | 16 | cg09821790 | 68283575 | rs9935025 | 68307447 | NFATC3 | NFATC3 | 0.0038 | -0.0201 | 0.0363 |
| 4 | 16 | cg09821790 | 68283575 | psy_rs55757091 | 68323654 | NFATC3 | NFATC3 | 0.002 | 0.0297 | 0.031 |
| 4 | 16 | cg09821790 | 68283575 | exm1251596 | 68344696 | NFATC3 | SMPD3 | 0.0028 | 0.0201 | 0.0329 |
| 4 | 16 | cg09821790 | 68283575 | rs13336173 | 68345691 | NFATC3 | SMPD3 | 0.0006 | 0.0233 | 0.0229 |
| 4 | 16 | cg09821790 | 68283575 | rs7189887 | 68353915 | NFATC3 | SMPD3 | 0.0066 | 0.0288 | 0.05 |
| 4 | 16 | cg09821790 | 68283575 | rs1868158 | 68398924 | NFATC3 | SMPD3 | 0.0021 | 0.0209 | 0.031 |
| 4 | 16 | cg09821790 | 68283575 | rs1122720 | 68412028 | NFATC3 | SMPD3 | 0.0008 | -0.0222 | 0.0229 |
| 4 | 16 | cg09821790 | 68283575 | rs8050499 | 68428326 | NFATC3 | SMPD3 | 0.0119 | 0.0229 | 0.0768 |
| 9 | 1 | cg20674738 | 26127400 | rs2786875 | 26061266 | PAQR7 | PAQR7 | 0.0001 | 0.0273 | 0.0993 |
| 9 | 1 | cg12709196 | 26289989 | rs2232648 | 26496455 | PAQR7 | CD52 | 0.001 | 0.0097 | 0.0993 |
| 9 | 1 | cg12709196 | 26289989 | exm34246 | 26582091 | PAQR7 | CD52 | 0.0018 | 0.0082 | 0.0993 |
| 9 | 8 | cg23130097 | 119208244 | rs4876852 | 119542307 | NA | NA | 0.0008 | 0.0153 | 0.0069 |
| 10 | 1 | cg27170383 | 37692359 | rs3762352 | 38156902 | MEAF6 | SNIP1 | 0.0005 | 0.0092 | 0.0873 |
| 10 | 1 | cg27170383 | 37692359 | rs11264087 | 38163387 | MEAF6 | SNIP1 | <0.0001 | 0.0123 | 0.0873 |

**Supp. Table 7.** Association between top response SNPs for dominance effect and topic score.

| SNP | Res_pval | Topic | Topic_pval | Coef | Chrom | Pos | Gene |
| --- | --- | --- | --- | --- | --- | --- | --- |
| rs6663620 | 1.82E-05 | NA | NA | NA | 1 | 90535344 | NA |
| rs12746015 | 7.57E-05 | 5 | 0.0803 | 0.1709 | 1 | 212636824 | NA |
| rs12746015 | 7.57E-05 | 9 | 0.0819 | -0.1708 | 1 | 212636824 | NA |
| rs11130670 | 8.72E-05 | 4 | 0.0488 | 0.1953 | 3 | 58658454 | NA |
| rs11130670 | 8.72E-05 | 9 | 0.0971 | -0.1649 | 3 | 58658454 | NA |
| exm-rs7901695 | 4.71E-05 | 5 | 0.0717 | 0.1721 | 10 | 114754088 | TCF7L2 |
| exm-rs4506565 | 4.71E-05 | 5 | 0.0717 | 0.1721 | 10 | 114756041 | TCF7L2 |
| rs2090852 | 4.55E-05 | NA | NA | NA | 12 | 51086931 | ATF1 |
| rs10783387 | 1.55E-05 | 5 | 0.0817 | 0.1673 | 12 | 51180143 | ATF1 |
| exm1002721 | 2.39E-05 | NA | NA | NA | 12 | 51203371 | ATF1 |
| exm2273492 | 8.50E-05 | NA | NA | NA | 12 | 51213765 | ATF1 |
| rs10774961 | 3.59E-05 | 4 | 0.0058 | -0.2671 | 12 | 109518133 | ACACB |
| rs12596798 | 7.68E-05 | 5 | 0.04 | 0.1972 | 16 | 8513277 | TMEM186 |
| rs257868 | 8.29E-05 | 4 | 0.0698 | -0.175 | 16 | 29412503 | SPN |
| psy_rs36025325 | 1.85E-05 | 10 | 0.097 | -0.1594 | 16 | 64075157 | NA |
| rs6085122 | 4.85E-05 | 4 | 0.0584 | 0.1822 | 20 | 5329782 | PROKR2 |
| rs6085122 | 4.85E-05 | 5 | 0.0108 | 0.2424 | 20 | 5329782 | PROKR2 |
| rs6085122 | 4.85E-05 | 9 | 0.0386 | -0.1988 | 20 | 5329782 | PROKR2 |
| rs6012442 | 9.25E-05 | 5 | 0.0157 | 0.2327 | 20 | 47105033 | ARFGEF2 |

